# Expanding the utility of transcriptome analysis for mutation detection in high-risk childhood precision oncology

**DOI:** 10.1101/2025.06.25.661445

**Authors:** Chelsea Mayoh, Paulette Barahona, Angela Lin, Lujing Cui, Pamela Ajuyah, Ann Altekoester, Loretta MS Lau, Dong-Anh Khuong-Quang, Patricia Sullivan, Akanksha Senapati, Sumanth Nagabushan, Ashleigh Sullivan, Natacha Omer, Andrew S Moore, Wayne Nicholls, Raelene Endersby, Nicholas G Gottardo, Geoffrey McCowage, Luciano Dalla Pozza, Jordan R Hansford, Seong-Lin Khaw, Paul J Wood, Toby N Trahair, Glenn M Marshall, David S Ziegler, Vanessa Tyrrell, Michelle Haber, Marie Wong, Paul G Ekert, Mark J Cowley

**Affiliations:** Children’s Cancer Institute, Lowy Cancer Research Centre, UNSW, Kensington, NSW, Australia; School of Clinical Medicine, UNSW Medicine & Health, UNSW Sydney, Kensington, NSW, Australia; University of New South Wales Centre for Childhood Cancer Research, UNSW, Sydney, NSW, Australia; Kids Cancer Centre, Sydney Children’s Hospital, Randwick, NSW, Australia; Children’s Cancer Centre, Royal Children’s Hospital, Parkville, Victoria, Australia; Murdoch Children’s Research Institute, Royal Children’s Hospital, Parkville, Victoria, Australia; Royal Manchester Children’s Hospital, Manchester, UK; Oncology Service, Children’s Health Queensland Hospital & Health Service, Brisbane, Queensland, Australia; Frazer Institute, Faculty of Medicine, The University of Queensland, Woolloongabba, Queensland, Australia; Child Health Research Centre, The University of Queensland, Brisbane, Queensland, Australia; School of Medicine, The University of Queensland, Brisbane, Queensland, Australia; Brain Tumour Research Program, The Kids Research Institute Australia, Nedlands, Western Australia, Australia; Centre for Child Health Research, University of Western Australia, Nedlands, Western Australia, Australia; Department of Paediatric and Adolescent Oncology and Haematology, Perth Children’s Hospital, Nedlands, Western Australia, Australia; Cancer Centre for Children, Children’s Hospital Westmead, Westmead, NSW, Australia; Michael Rice Centre for Haematology and Oncology, Women’s and Children’s Hospital, South Australia Health and Medical Research Institute, Adelaide, South Australia, Australia; South Australia ImmunoGENomics Cancer Institute, University of Adelaide, Adelaide, South Australia, Australia; Monash Children’s Hospital, Melbourne, Victoria, Australia; Centre for Cancer Research, Hudson Institute of Medical Research, Clayton, Victoria, Australia; Department of Paediatrics, School of Clinical Sciences at Monash Health, Monash University, Clayton, Victoria, Australia; Cancer Immunology Program, Peter MacCallum Cancer Centre, Parkville, Victoria, Australia; The Sir Peter MacCallum Department of Oncology, University of Melbourne, Parkville, Victoria, Australia

## Abstract

In precision oncology, whole transcriptome sequencing (RNA-seq) excels at identifying oncogenic fusions. Here, using a cohort of 477 high-risk paediatric tumours, we demonstrate that RNA-seq can identify all mutation classes found previously using whole genome sequencing (WGS) and provides additional functional insights into their pathogenicity. By incorporating reference-guided fusion, and reference-free structural variant (SV) detection algorithms with RNA abundance assessment, RNA-seq identified 96% of SVs and resolved 33 complex SVs that WGS failed to identify. Furthermore, RNA-seq identified 92% of all single nucleotide variants and small insertions and deletions. Importantly RNA-seq informed the pathogenicity assessment in 22% of variants through identification of allele specific expression or the functional consequence of splice-altering variants. The utility of RNA-seq extends beyond fusion identification to the interpretation of mutation pathogenicity and the discovery of important mutations that would otherwise go undetected. We propose that RNA-seq is an indispensable companion to WGS in precision medicine.

## Introduction

In cancer precision medicine programs, transcriptome sequencing (RNA-seq) is a critical tool for guiding patient diagnosis and aiding treatment decisions. RNA-seq provides insight into the functional consequences of mutations, providing evidence that is essential to interpreting pathogenicity. Currently, existing RNA-seq approaches in precision oncology are primarily used for detection of oncogenic fusions^1^, leaving a wealth of additional information underutilised. This highlights the need to expand these approaches to fully leverage the broader potential of RNA-seq in precision oncology.

Paediatric cancers have a significantly lower mutation burden than adult cancers and are frequently driven by structural variants (SV)^2–4^. These may be large deletions, duplications, insertions, inversions and translocations, or smaller intragenic SVs only affecting parts of a gene^5^. The accurate identification of expressed SVs, their reading frame and retained protein domains, are imperative for accurate identification of cancer drivers and identifying personalised treatment recommendations (i.e. NTRK fusion with a retained tyrosine kinase domain indicative of response to Larotrectinib)^6^.

Reference-guided fusion detection algorithms align RNA-seq reads to a reference genome with known exons and splice junctions^7–9^, making them effective at detecting fusions with well-characterised genes and canonical splice sites. However, they have limitations in their ability to detect novel SVs, particularly those that create non-canonical splice junctions (e.g. *RB1* intragenic deletions). A complementary approach is to use a reference-free RNA-seq method which assembles RNA-seq reads into transcripts without prior knowledge of exon boundaries or splice junctions^10^. This enables the accurate detection of novel isoforms and/or splice junctions that result in aberrant gene splicing, which is critical for identifying clinically actionable targets and understanding the biological mechanisms driving tumorigenesis in high-risk paediatric cancers.

Single-nucleotide variants (SNVs) and small insertions and deletions (Indels, <50 base pairs) are also important drivers of high-risk paediatric cancer^2,3,11^, which can be identified from RNA-seq data, yet are underutilised in precision oncology. Furthermore, RNA-seq can reveal the functional consequences of these mutations, aiding in their interpretation^12^. RNA-seq has the potential to identify the abundance of transcribed mutations, identify preferential expression of one allele (i.e. allele specific expression), confirm nonsense mediated decay resulting from stop-gain mutations, and determine the transcribed consequence of splicing mutations. Thus, extending the utility of RNA-seq to the analysis and interpretation of DNA mutations, provides functional insight not possible from DNA sequencing alone.

The ZERO Childhood Cancer program (ZERO) is Australia’s national precision medicine program for all paediatric cancers^11,13^. The published description of the first 247 high-risk patients showed that the ZERO analytical platform, which includes germline and tumour WGS and tumour RNA-seq profiling identified a pathogenic or likely-pathogenic aberration in 94% of these patients^11^, higher than other reported paediatric precision medicine programs^14–16^. In ZERO, RNA-seq was key to the higher rate of detection of pathogenic variants, but 42% of the variants were only detected by WGS. Here, we present the strategies developed since then to enhance the utility of RNA-seq in precision oncology and improve the detection of SVs, SNVs, and Indels. We demonstrate how RNA-seq data can resolve the functional consequences of DNA mutations and aid in the accurate interpretation of variant pathogenicity. In addition, we highlight how RNA-seq resolves the transcriptional consequences of complex structural rearrangements to identify expressed oncogenic drivers. To facilitate the adoption of our findings, we developed *Carbonite*, an open-source scalable RNA-seq analysis pipeline for cancer precision medicine. We propose that RNA-seq has broad applications in tumour genomic analyses and, when combined with WGS, is an indispensable tool in the characterisation of cancer genomes and the identification of targeted therapies.

## Results

### Establishment of a comprehensive paediatric high-risk cohort to identify pathogenic variants in transcriptome sequencing

We analysed data from the first 477 consecutively enrolled high-risk paediatric cancer patients in the ZERO Childhood Cancer precision medicine program, enrolled between June 2015 and May 2021, that had matched WGS and RNA-seq data from the same tumour biopsy. Our previous studies^11,13^ have reported on WGS-based mutation findings in 228 then 319 of these patients; this study includes those cohorts and adds data from 158 newly sequenced patients. These included 182 tumours of the central nervous system (CNS), 120 sarcomas, 64 solid tumours (ST), 36 neuroblastomas (NB), and 75 haematological malignancies (HM) (Extended Data Figure 1A). Every sample was manually curated to identify clinically relevant pathogenic and likely pathogenic variants, which we refer to as “reportable”, and presented to a multi-disciplinary molecular tumour board to aid clinical management as previously described^11,13^. Here, we focused only on the reported structural variants (SV), SNVs or Indels. Reportable aberrations were identified independently between DNA and RNA with the output from each platform manually integrated. This manually integrated WGS and RNA-seq pipeline identified a total of 755 reportable aberrations with 218 SVs and 537 SNVs/Indels (Extended Data Figure 1B).

### Identification of driver structural variants using reference-guided fusion calling in RNA-seq

In paediatric cancer, it is known that all types of SVs including translocations, deletions, duplications, inversions and insertions drive disease^17,18^. While WGS usually identifies the exact breakpoints of SVs, their functional consequences are best observed using RNA-seq. Typically multiple fusion detection algorithms are used in RNA-seq analysis pipelines to balance the strengths and limitations of individual algorithms, which largely derive from differences in alignment-based strategies and filtering criteria^7–9^. In ZERO, the RNA-seq analytical pipeline processes the data through three reference-guided RNA-seq (rgRNA-seq) algorithms Arriba, STAR-Fusion and JAFFA^7–9^. Of the 218 reportable SVs 69% were identified using at least one rgRNA-seq method (Figure 1A). Seventy percent (105/150) of these SVs were identified by all three callers, and of these 105, Arriba identified 95%, STAR-Fusion 81.2% and JAFFA 80.5% (Extended Data Figure 2A). Arriba excelled through its identification of both gene fusions and other SVs such as an *HGF* internal tandem duplication (ITD) in a malignant rhabdoid tumour and an *IKZF1* inversion in a Ph-like pre-cursor B acute lymphoblastic leukaemia (ALL) (Extended Data Figure 2B–C); both were missed by STAR-Fusion and JAFFA. However, other key driver SVs including a *P2RY8::CRLF2* gene fusion in a Ph-like ALL, were flagged by Arriba as potential false positives, reinforcing the importance of using multiple fusion callers.

**Figure 1.**
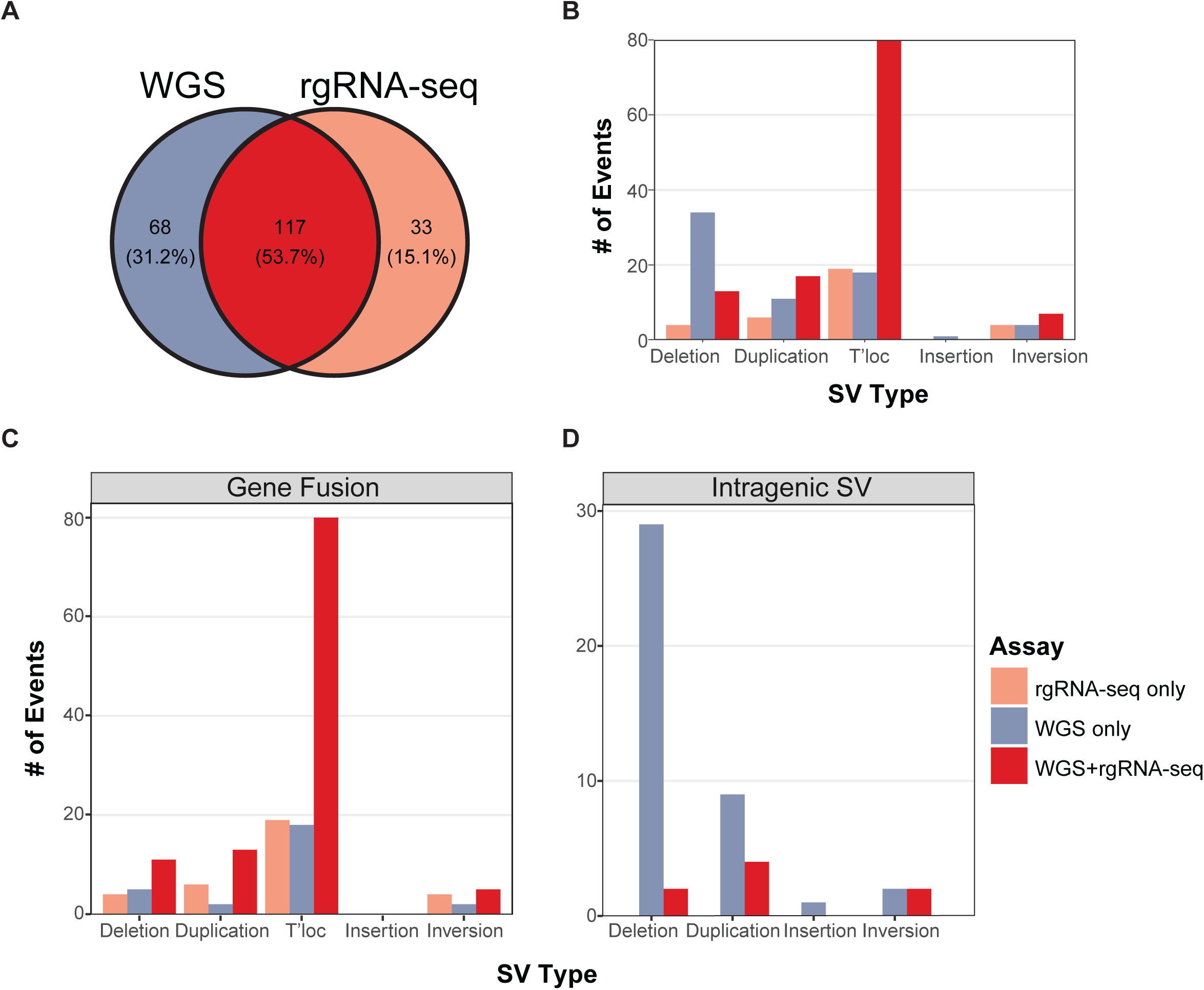
Comparison of structural variants identified by WGS and reference-guided RNA-seq. (A) Venn diagram showing the overlap of 218 structural variants (SV) detected by whole genome sequencing (WGS) and three reference-guided RNA-seq (rgRNA-seq) methods. The number of total SVs (**B**), gene fusions (**C**), and intragenic SVs (**D**) identified by rgRNA-seq (peach), WGS (purple) or both (red), stratified by SV type: deletion, duplication, translocation (T’loc), insertion and inversion.

To understand why rgRNA-seq missed 31% of all reportable SVs (Figure 1A), we examined detection rates across different SV types. Unsurprisingly, rgRNA-seq excelled at identifying gene fusions caused by translocations, where DNA from two chromosomes are joined, successfully detecting 85% (99/117) of these events (Figure 1B). Detection was moderate for inversions and duplications, with 73% (11/15) and 68% (23/34) identified, respectively (Figure 1B). In contrast, performance was poor for deletions, with only 33% (17/51) detected (Figure 1B), and the single insertion event was missed. To better understand these discrepancies, we grouped SVs into two broader categories: gene fusions (formed from 2 genes), or intragenic SVs (occurring within a single gene). This revealed that rgRNA-seq identified 84% (142/169) of all gene fusions (Figure 1C), but only 16% (8/49) of intragenic SVs (Figure 1D). These findings highlight a considerable opportunity to improve RNA-seq pipelines for more comprehensive detection of all driver SVs.

### RNA-seq pipeline incorporating RNA gene abundance and a reference-free analysis identifies structural variants previously seen only in WGS

To understand why our rgRNA-seq-based SV detection pipeline failed to identify 16% (27/169) of gene fusions, we examined the characteristics of these missed fusions in more detail (Figure 2A). Of the 27 missed fusions, 12 were predicted by WGS to activate oncogenes, while 15 disrupted tumour suppressor genes (TSGs). Strikingly, 67% (8/12) of the missed oncogenic SVs involved breakpoints in non-coding regions (Figure 2A), consistent with enhancer or promoter hijacking mechanisms predicted to drive oncogene activation^19^. Of particular interest were two non-coding enhancer hijacking SV events affecting *MYC* and three promoter hijacking SV events affecting *TERT* (Figure 2B–C). We reasoned that if these events were functional this would result in increased expression of these oncogenes. To examine this, we performed z-score analysis to detect outlier expression of *MYC* or *TERT* compared to wild-type (WT) controls lacking genetic alterations in these genes. All five non-coding SVs resulted in increased expression of *MYC* or *TERT*, providing functional evidence supporting the activation of these oncogenes, thus the pathogenicity of the SV (Figure 2D). While most hijacking events involved intergenic breakpoints, we also identified five events occurring within the transcribed *MYC* sequence resulting in detection in rgRNA-seq. An example being the complex *MYC::IGH* rearrangement with previously defined breakpoint architecture^20,21^ that generates chimeric reads within the transcribed region, and another example the previously reported *PVT1::MYC* fusion^22^ where the first exon of *MYC* is not transcribed, due to it lacking a 5’ splice site (Extended Data Figure 2D–E).

**Figure 2.**
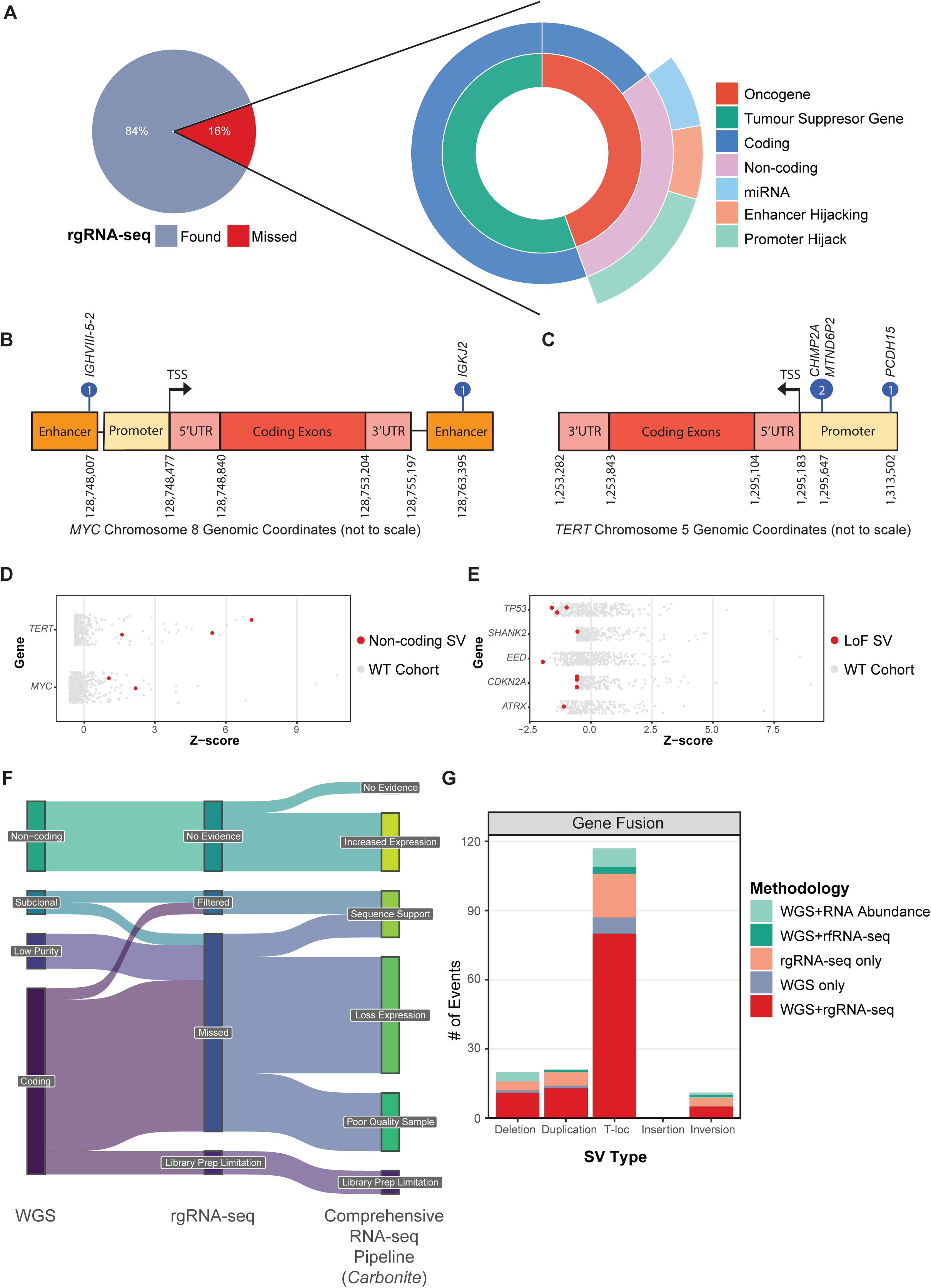
Outlier gene abundance and reference-free RNA-seq identify coding and non-coding gene fusions. (A) The proportion of SVs detected or missed using rgRNA-seq methods. Missed gene fusions were further classified by gene type (oncogene or tumour suppressor gene; inner ring), breakpoint location (coding or non-coding; middle ring), and non-coding category (miRNA cluster, putative enhancer or promoter hijacking events; outer ring). Schematic of non-coding enhancer hijacking SVs involving *MYC* **(B)** and promoter hijacking SVs involving *TERT* **(C).** Relative RNA abundance (z-score) of oncogenes affected by non-coding SVs (red; **D**) and tumour suppressor genes affected by loss of function SVs (red; **E**), compared to wildtype (WT) controls (grey). **(F)** Sankey diagram of gene fusions initially detected only by WGS (left), reasons for their absence from rgRNA-seq fusion detection algorithms (middle) and the supporting evidence from *Carbonite*, or the rationale for non-detection (right). **(G)** The number of gene fusions stratified by SV type: deletion, duplication, translocation (T’loc), insertion and inversion, and their detection method: rgRNA-seq only (peach), WGS only (purple), both (red), WGS and reference-free RNA-seq (dark green), or WGS with RNA abundance support (teal).

In contrast, coding fusions involving TSGs often result in truncation and loss of function, but typically show no supporting read evidence in RNA-seq due to reduced or absent gene expression. Here, we identified such events affecting *TP53*, *SHANK2*, *EED*, *CDKN2A*, and *ATRX*. Using the same z-score analysis, we confirmed lower expression of the affected TSGs compared to WT controls, providing orthogonal validation of the damaging nature of the SV (Figure 2E). These findings demonstrate that integrating RNA abundance analysis into a RNA-seq pipeline enhances the interpretation of both non-coding SVs affecting oncogenes and loss-of-function SVs in TSGs. This is further supported by our previous work, which demonstrated that transcript abundance can serve as a valid therapeutic biomarker and correlate with treatment response^13^.

There were a further two non-coding gene fusions that went undetected, both being rearrangements of the C19MC miRNA cluster which is not captured with the polyA mRNA sequencing protocol used (see methods), reflecting a library prep limitation (Figure 2F). Taken together, we have identified the reason and shown either direct or supporting evidence for 18/27 (67%) of the missed SVs. To detect the remaining nine missed coding gene fusions, we hypothesised that a reference-free RNA-seq (rfRNA-seq) approach, such as MINTIE, would enable their detection. MINTIE is an algorithm that performs *de novo* assembly and was developed to identify cryptic SVs and novel splicing events^23^. The challenge of distinguishing true calls using MINTIE from false positive calls arising from RNA-seq artefacts or intron sequences was substantially mitigated by restricting the calls to those made from the matched WGS data. In this way, four (44%) coding gene fusions were recovered (Figure 2F). All had low but definite supporting reads, showing that even where there are few reads, rfRNA-seq can identify gene fusion events. The remaining five coding gene fusions that rgRNA-seq missed were from samples with degraded RNA (i.e. RIN <5). Thus, by expanding beyond a RNA-seq pipeline using reference-guided fusion detection algorithms and incorporating RNA abundance and a reference-free approach, we were able to provide evidence for 95% (161/169) of gene fusions and identified the reason rgRNA-seq could not detect in all remaining cases where no evidence had been observed (Figure 2F-G, Extended Data Figure 3A).

### Incorporating reference-free analysis and RNA gene abundance improves intragenic SV detection

We mapped the anticipated transcriptional outputs of intragenic SV events to guide our development of more accurate detection of these events in RNA-seq data (Figure 3A). Intragenic breakpoints linking non-adjacent exons may result in aberrant isoforms (Figure 3A). Inversions and deletions that disrupt the transcriptional start site (TSS) may abolish gene transcription, unless an alternative downstream TSS is present, resulting in expression of a shorter isoform. Whereas intragenic duplications of exon 1 will not alter the mature splice mRNA sequence, nor should they affect abundance of the resulting transcript (Figure 3A). Using this conceptual framework, we applied the rfRNA-seq algorithm MINTIE, which can also identify the functional consequence of intragenic SVs through the creation of novel splice junctions. MINTIE recovered 74% (23/31) of the intragenic deletion and inversion events identified in WGS but missed by rgRNA-seq, with all classified as aberrant splicing (Figure 3B). For example, an intragenic deletion of exons 2-24 in *ATRX* in an adrenocortical carcinoma resulted in a novel splice junction between exons 1–25 (Figure 3C). Further examples of using aberrant splicing detection to identify intragenic deletions in *PMS2* and *CTNNB1* are shown in Extended Data Figure 4A–B. rfRNA-seq also identified 89% (8/9) of oncogenic ITDs missed by rgRNA-seq, including driver events in *BCOR*, *FGFR1*, and *MYC* (Extended Data Figure 3B). The duplication not recovered by rfRNA-seq was a *FGFR1* ITD; however, following manual review in the integrated genomics viewer (IGV), we identified a clear increase in the number of reads spanning the duplicated exons with supporting soft-clipped reads. The only genomic insertion in our dataset was detected by rfRNA-seq and involved the insertion of 17 nucleotides from chromosome 3 into intron 36 of *NF1* in an H3 K27-altered diffuse midline glioma. This created a toxic exon that triggered aberrant splicing and led to loss of function of the tumour suppressor gene (Extended Data Figure 4C). Thus, by incorporating a rfRNA-seq approach into our RNA-seq pipeline, we recovered 76% of the missed intragenic SVs identified by WGS (Figure 3B).

**Figure 3.**
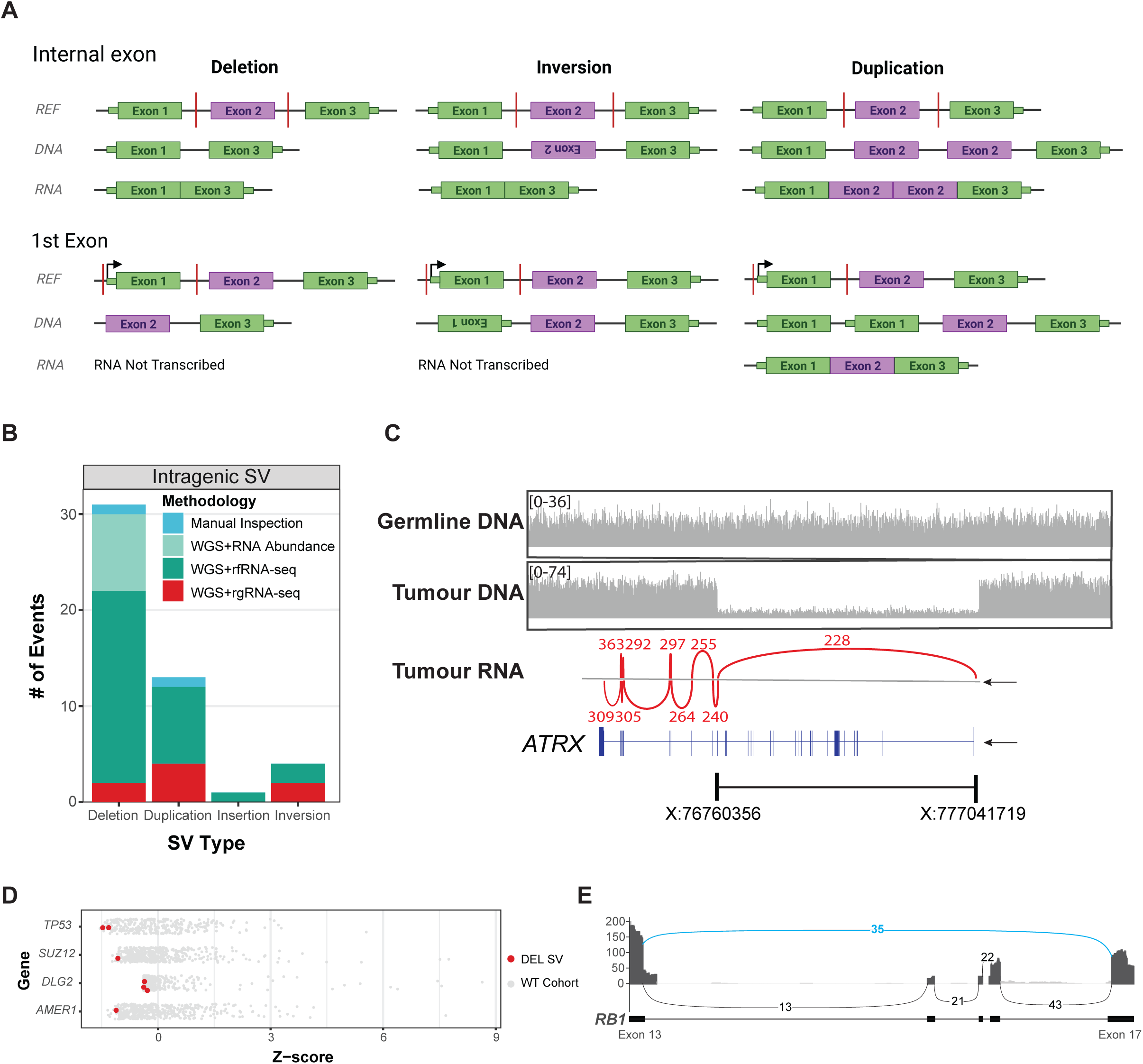
Reference-free RNA-seq and gene abundance reveal intragenic structural variants. (A) Schematic illustrating the functional consequences of intragenic SVs affecting internal (top) or first exons (bottom), with reference sequence, breakpoint positions, and resulting effects in tumour DNA and RNA for deletions (left), inversions (middle), and duplications (right). (**B**) The number of intragenic SVs detected by WGS and rgRNA-seq (red), WGS and reference-free RNA-seq (dark green), WGS with RNA abundance changes (teal), or manual inspection using IGV (blue), stratified by SV type: deletion, duplication, insertion, and inversion. (**C**) Intragenic deletion of *ATRX*, showing germline DNA coverage (wild-type, top), tumour DNA coverage (biallelic deletion, middle), tumour RNA sashimi plot with splice-junction support, exon structure and transcriptional direction, and genomic coordinates (hg19) of the deletion (bottom). (**D**) Relative RNA abundance (z-score) of tumour samples (red) compared to wild-type controls (grey), for SVs with 5’ breakpoints upstream of the transcriptional start site that disrupt at least the first exon of tumour suppressor genes. (**E**) Sashimi plot of tumour RNA from a sample with an intragenic deletion of exons 14–16 in *RB1*, showing exon skipping and a novel splice junction from exons 13 to 17 (blue).

For the nine remaining undetected intragenic SVs, the 5’ breakpoint occurred upstream of the TSS. In seven cases, these manifested as a loss of gene expression, as evidenced by having low expression (z-score ≤-1.5; Figure 3D). An *IKZF1* intragenic deletion (spanning from chr7:50345518-50440833) knocked out the TSS of the canonical isoform and resulted in a truncated alternate isoform and a non-functioning protein. Another intragenic deletion in *RB1* was not detected by our pipeline but was clearly evident in IGV as an aberrant splice junction (Figure 3E). Our data shows that incorporating additional modalities into an RNA-seq pipeline, notably reference-free algorithms that incorporate detection of aberrant splicing, and gene abundance measures, greatly improve SV detection to 96% (210/218) of all reportable SVs (Extended Data Figure 3).

### RNA-seq resolves expressed gene fusions arising from complex structural rearrangements and difficult to map regions of the genome

We next explored the 33 gene fusions that were exclusively identified from rgRNA-seq but not detected in WGS. There were three main classes of events: gene fusions that arose from multi-hop rearrangements that involved more than two contiguous genes in the DNA, SVs involving gene paralogs, and complex SVs arising from genome shattering events (Figure 4A). An example of a multi-hop event shown in Figure 4B–D involved *EML4*, *ITM2C*, *CAB39*, *BANF1P3* and *ALK* on chromosome 2 in a spindle cell tumour. rgRNA-seq identified a high-confidence, in-frame *EML4::ALK* fusion that included an intact tyrosine kinase domain (Figure 4C), a clinically important finding since this lesion may be targeted with ALK inhibitors. Visualization of the WGS data, together with the retrospective application of PURPLE, GRIDSS2 and LINX^24^ algorithms into the WGS pipeline, recovered this event by “chaining” the breakpoints together (Figure 4D). *CIC::DUX4* fusions are key drivers of CIC-rearranged sarcomas, but their detection is complicated by the presence of multiple DUX4 paralogs present on either chromosome 4 (Figure 4E) or chromosome 10. These paralogs also play a role in DUX4-rearranged B-cell acute lymphoblastic leukaemias. In this cohort, we identified two DUX4 fusions that were missed by WGS, likely due to the STAR aligner’s improved handling of read alignment in repetitive regions of the genome^25^. This principle also applies to the diagnostic synovial sarcoma fusions where *SS18* fuses to either the *SSX1* or *SSX2* genes which are paralogs (Extended Data Figure 3A). In the genome shattering class, rgRNA-seq resolved the transcriptional consequence of this event to identify an in-frame *KANK1::NTRK2* fusion of a typical structure, with a 5’ protein-protein interaction domain and a 3’ tyrosine kinase domain (Figure 4F). WGS revealed that a complex SV event had occurred, with several chained breakpoints (Figure 4G). Regardless of the underlying genomic complexity, the spliced mRNA sequence revealed a novel splice junction linking *KANK1* to *NTRK2*, resulting in an activating fusion of a clinically targetable oncogene. When we reanalysed WGS data from these samples using tools specifically designed to phase reads from complex genomic rearrangements^24^, 28 of the 33 gene fusion events initially detected only by rgRNA-seq were also detected (Extended Data Figure 3A), albeit still requiring the expert interpretation of less-familiar LINX visualizations. A further five events were not detected by WGS (Extended Data Figure 3A). Taken together, rgRNA-seq identified an additional 15% (33/218) of clinically relevant and in some instances, therapeutically targetable SVs that WGS SV algorithms either failed to detect or was unable to resolve.

**Figure 4.**
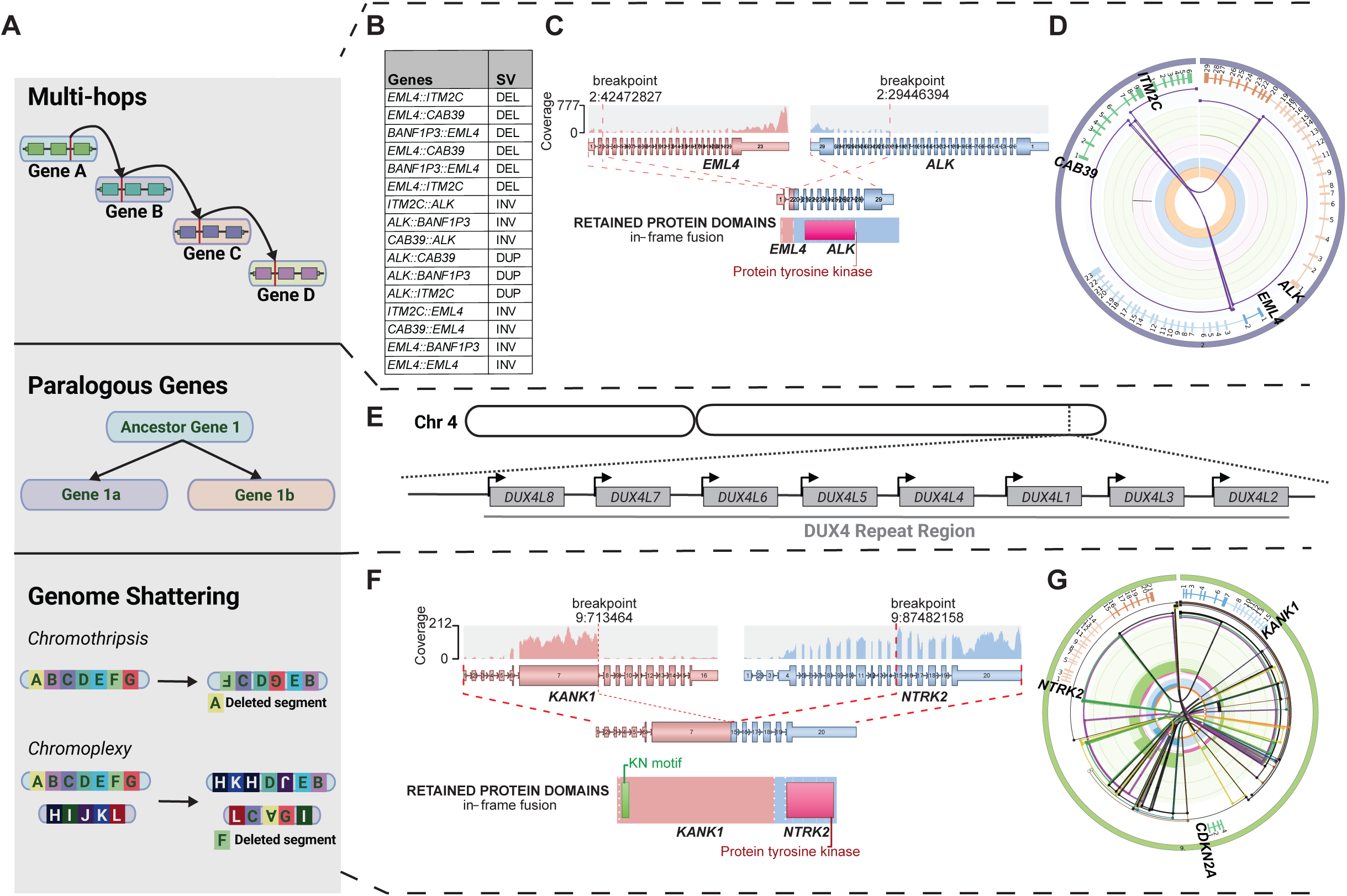
rgRNA-seq resolves complex and paralogous SVs missed by WGS. (A) Thirty-three SVs undetected by WGS were identified using rgRNAseq, falling into three major categories: gene fusions arising from multi-hop rearrangements (top), gene fusions involving gene paralogs (middle), and complex SVs, such as chromothripsis or chromoplexy (bottom). **(B)** Tabular output from WGS showing the breakpoints and gene fusion annotations from an example multi-hop SV event. **(C)** Arriba identified an in-frame *EML4::ALK* fusion retaining the protein tyrosine kinase domain. **(D)** The retrospective use of LINX reconstructed the multi-hop rearrangement linking *EML4* and *ALK* by chaining DNA breakpoints. **(E)** Schematic of the *DUX4* repeat array region on chromosome 4, illustrating multiple loci that impair DNA read alignment. **(F)** Arriba identified a *KANK1::NTRK2* fusion showing the transcribed exons, breakpoint locations, expression coverage, and retained in-frame protein domains. **(G)** LINX plot of a complex genome shattering event involving the *KANK1::NTRK2* SV, initially missed by WGS.

### RNA variant allele frequency enhances pathogenicity assessment of putative driver SNVs and Indels

SNV and Indel variants encode many critical driver events and understanding how they are expressed provides valuable insights into their functional impact. We developed an RNA-seq-based SNV/Indel analysis pipeline and assessed the contribution of this to the pathogenicity assessment of putative driver alterations. WGS identified 537 reportable SNVs/Indels, of which 492 (91.6%) were also detected using RNA-seq-based variant calling (Figure 5A, Extended Data Figure 1B). Among these, 81% (397/492) showed concordant VAFs between WGS and RNA-seq, indicative of heterozygous or homozygous expression (Figure 5B – labelled “validation”, Supplementary Table 1). Importantly, RNA-seq provided additional insight into the pathogenicity of a variant, revealing allele specific expression (ASE) in 70 (13%) somatic variants, where heterozygous variants in the genome were expressed exclusively as the mutant allele in RNA (Figure 5B - labelled “het2hom”). These events are particularly important in TSGs, where loss of expression of the WT allele indicates complete functional inactivation. ASE was most prevalent in *TP53*, but also frequently observed in *SMARCA4*, *NF1* and *CDKN2A*, all key TSGs in paediatric high-risk cancers (Supplementary Table 1). In contrast, RNA-seq showed no expression of 24 WGS-identified SNVs/Indels, with only the WT allele represented in the transcriptome. Ten of these were Indels, but this was unlikely due to systematic technical limitations in Indel detection, as 89% (114/128) of WGS-reported Indels were successfully detected in RNA-seq (Figure 5C). A further 21 SNVs/Indels (3.9%) were not present in RNA-seq due to insufficient coverage of either the relevant exon or gene (Figure 5A, C, Supplementary Table 1). Notably, seven (16%) of these were stop-gain mutations and their lack of expression suggests degradation via nonsense-mediated decay, further supporting their pathogenicity (Figure 5A-C). Together, these findings demonstrate that integrating RNA-seq VAF and expression data provides orthogonal evidence for variant interpretation, enhancing confidence in both gain- and loss-of-function events and refining the classification of driver mutations.

**Figure 5.**
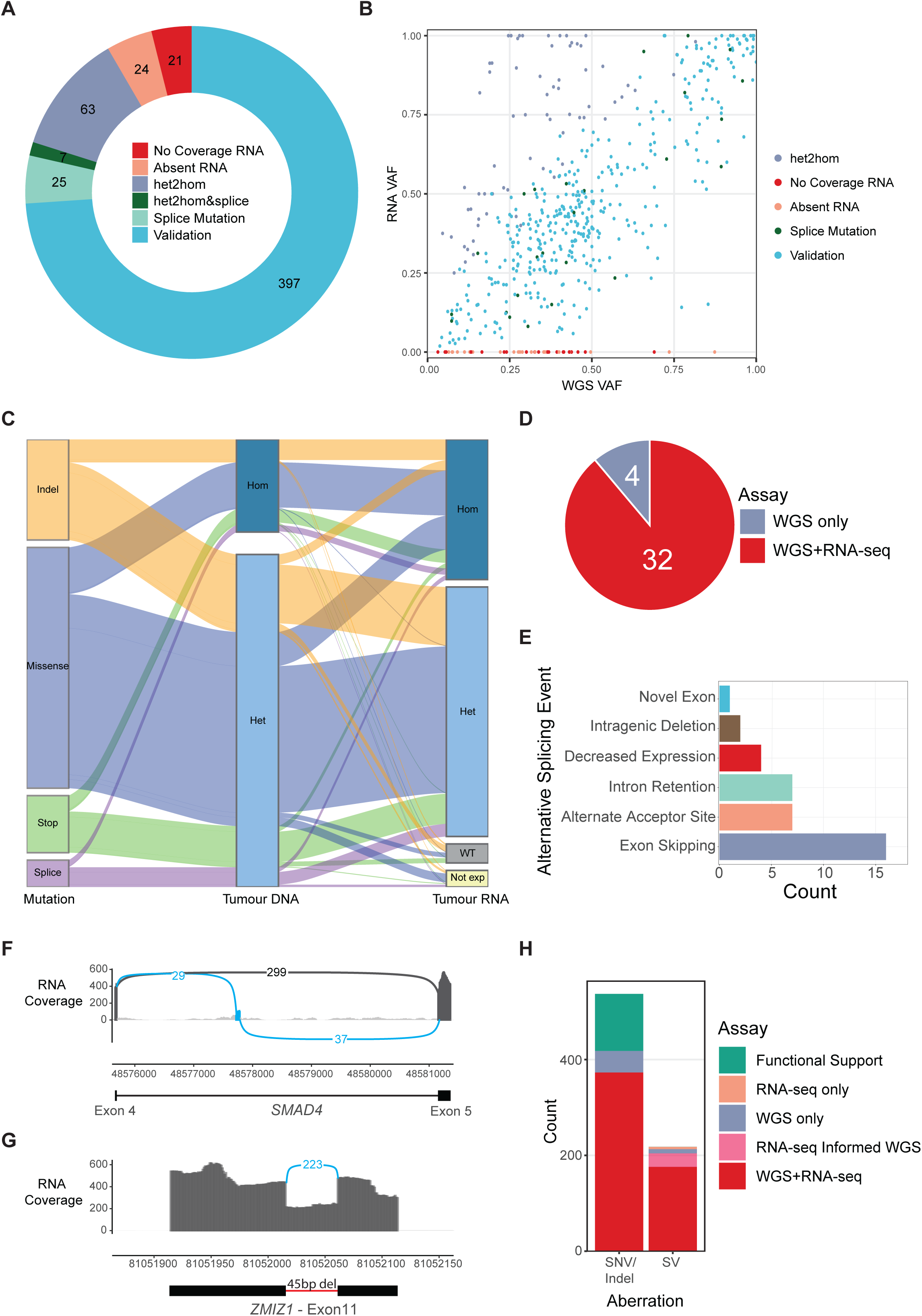
RNA-seq orthogonally validates SNV/Indel mutations and reveals allele-specific expression and aberrant splicing. (A) Summary of 537 SNV/Indel mutations identified in DNA and assessed in RNA-seq. Variants were classified as orthogonally validated (blue) when the variant allele was observed in both WGS and RNA-seq at similar zygosity, no RNA coverage (red), allele-specific expression of either the wild-type allele (peach), or the mutated allele (purple), splice site mutation affecting just one (teal) or both alleles (dark green). **(B)** Scatterplot of variant allele frequencies (VAF) from WGS (x-axis) and RNA-seq (y-axis), coloured as in A. **(C)** Alluvial plot showing mutation type: Indel (orange), missense (purple), stop-gain (green), and splice (light purple); tumour DNA variant zygosity from WGS (*heterozygous*, light blue, or *homozygous*, dark blue); and RNA zygosity (heterozygous, homozygous or wildtype [grey]) or not expressed (light yellow). **(D)** The number of splice-site mutations with RNA-seq evidence of aberrant splicing (red) or no RNA coverage (purple). **(E)** Alternative splicing events detected in RNA-seq, classified as exon skipping (purple), alternate acceptor site (peach), intron retention (teal), intragenic deletion (dark brown), novel exon (blue), or decreased gene expression (red). Sashimi plots highlight an example of **(F)** a novel toxic exon resulting from a deep intronic deletion disrupting the splice acceptor site in *SMAD4* intron 4 and **(G)** a 45-base pair (bp) intragenic deletion in exon 11 of *ZMIZ1* identified by *RNAindel*. **(H)** Following enhanced mutation detection by *Carbonite*, SNV/Indels and SVs were classified by detection platform and supporting evidence: WGS only (purple), RNA-seq only (peach), both (red), initially detected in RNA-seq and later confirmed in WGS via LINX (pink), or functionally supported by RNA-seq (green) including allele-specific expression or aberrant splicing indicative of splice-site disruption.

Determining the functional consequence of putative splice-altering variants using DNA sequencing alone remains challenging, even with state of the art *in silico* prediction tools^26,27^. In contrast, RNA-seq provides direct evidence of the functional consequence of such variants. Among the 36 reportable splice-altering mutations identified in our dataset, aberrant splicing was detected in 32 cases (89%) using RNA-seq (Figure 5D). Of these, fifty percent (16/32) resulted in exon skipping and 22% (14) involved intron retention or the use of alternative splice sites (Figure 5E). Uncommon splice alterations included the formation of a novel 62 base-pair exon in *SMAD4* (Figure 5F), a 45 base-pair deletion in *ZMIZ1* (Figure 5G) and a 30 base-pair deletion in *MYC*. Notably, seven of the splice-altering variants were heterozygous at the DNA level but showed exclusive expression of the aberrant isoform in RNA-seq, indicating ASE and functional homozygosity (Figure 5C, Supplementary Table 1). Thus, RNA-seq not only revealed the functional consequences of these splice variants but, when integrated with ASE analysis, provided strong evidence for complete loss of normal splicing in key TSGs such as *NF1*, *TP53* and *WT1* (Supplementary Table 1).

Interestingly, four splice-altering mutations showed no detectable impact on isoform structure but were associated with reduced gene expression. In these cases, the lack of transcript-level evidence made it difficult to rule out a splicing effect (Supplementary Table 1), and accordingly, we classified these variants as potentially splice-altering. Overall, through the combination of ASE analysis and the direct confirmation of aberrant splicing, RNA-seq contributed new functional insights into the pathogenicity of 22% of all SNVs/Indels in our cohort.

## Discussion

In precision oncology, a major goal is to identify all clinically relevant driver alterations in every tumour, particularly those that inform treatment decisions, aid diagnosis or provide prognostic insights. Our findings demonstrate that tumour RNA-seq analysis can identify more clinically significant information than its traditional use for gene fusion detection alone. In this study, we demonstrate how a comprehensive RNA-seq analysis, integrated with WGS, can detect and confirm intergenic and intragenic SVs, reveal the consequences of splice-altering mutations, validate the allele specific expression of SNVs and Indels, and resolve the functional impact of complex genomic events such as chromothripsis. Overall, 96% of reportable SVs, SNVs and Indels identified by WGS were also identified by *Carbonite*, our integrative RNA-seq pipeline, effectively closing the ‘mutation detection gap’ from our earlier work^11^. Moreover, the *Carbonite* pipeline provided additional functional evidence for 22% of SNVs/Indels and 15% of SVs, that were not captured by WGS analysis alone (Figure 5H).

Large-scale precision medicine programs in paediatric cancer, such as ZERO, have demonstrated that WGS detects more clinically relevant aberrations than whole exome sequencing (WES) or targeted panel sequencing^11,28^. While combining WGS and WES can help validate the detection of specific variants^29^, the high accuracy of both methods raises questions about the necessity, and cost, of this orthogonal validation. Our findings show that RNA-seq not only serves this orthogonal validation role but, more importantly, provides critical insights into the functional and pathogenic consequences of variants. The examples presented here, including the resolution of complex genomic events into targetable fusions, detection of allele specific expression of mutated TSGs, and confirmation of aberrant splicing in cancer genes, underscore the substantial added value of RNA-seq in refining genomic interpretation.

A key enhancement in *Carbonite* was deploying reference-free SV-detection algorithms, which helped overcome alignment biases inherent to reference-guided approaches^7–9,23^. The reference-free approach excels at detecting novel splice junctions and intragenic SVs. In some cases, it also accurately identifies breakpoints when fewer supporting reads are present, such as in low tumour purity samples, which if left undetected, would otherwise result in needing an additional tumour biopsy. However, a notable limitation of reference-free approaches is their tendency to identify large amounts of noise. While this can be mitigated using tumour-type matched controls, in our experience this can inadvertently exclude key driver events, such as the diagnostic *PAX3::FOXO1* fusion. To address this, our approach omits control-based filtering and instead is guided by SVs identified from WGS. The development of sophisticated filtration algorithms or set of cancer type specific controls may reduce the need for WGS, strengthening the utility of an RNA-seq only assay for mutation detection.

We found that rgRNA-seq methods were able to identify important driver fusions from genes that have paralogs, including *DUX4* and *SS18*, that were missed by WGS. Our WGS analysis pipeline uses BWA-mem^30^, which marks multi-mapping reads with a mapping quality of 0 and randomly picks one location as the primary alignment. As most variant detection tools ignore reads with poor mapping quality, this supporting evidence is missed. In contrast, the STAR aligner has been demonstrated to outperform other rgRNA-seq alignment tools in repetitive regions^25^. We attribute the enhanced performance of fusion detection in rgRNA-seq in repetitive regions to both this enhanced read alignment and having additional supporting read evidence from highly expressed gene fusions. Moreover, rgRNA-seq proved valuable in resolving complex SVs that can obscure actionable driver lesions, such as those arising from chromothripsis or chromoplexy. Our identification of an *NTRK2* fusion arising in this way (see Figure 4) is to our knowledge, a first. This underscores the importance of interpreting complex SVs through their transcribed consequences, and we argue that RNA-seq is the most facile way to achieve this. A critical aspect of assessing the pathogenicity of novel fusion events, especially those involving tyrosine kinases^6,31–33^, is verifying that the resulting transcript retains the functional domains that drive oncogenicity. For example, receptor tyrosine kinase fusions typically require an intact 5’ protein-protein interaction domain and a 3’ kinase domain that are in-frame. Our study demonstrates that RNA-seq enables accurate interpretation of retained protein domains, providing the additional functional insights into SV oncogenicity. Additionally, RNA-seq confirmed loss-of-function events in TSGs like *NF1* and *SMARCB1,* which may inform therapeutic decisions, such as the use of MEK-inhibitors or EZH2-inhibitors respectively, both of which have shown objective clinical benefit^13^. Taken together, our results support the essential role of RNA-seq in identifying diagnostically and therapeutically relevant alterations within complex genomic backgrounds, providing critical insights that may be missed by WGS alone.

Assessing ASE provides critical insights into the functional impact of SNVs and Indels. This is particularly relevant for TSGs, where heterozygous somatic mutations are functionally homozygous if the mutant allele is exclusively expressed in the tumour. In our cohort, this was most frequently observed in *TP53*. Although our data does not clarify the mechanisms by which the wildtype allele is silenced, future studies could integrate mutation data with DNA methylation data to investigate the roles of promoter hypermethylation. In addition, albeit less common, an activating mutation even if therapeutically targetable, is clinically irrelevant if not expressed. This phenomenon was observed in just 4% of cases and may explain if a patient has unexpectedly poor response to a targeted therapy.

We have shown that *Carbonite*, which integrates reference-guided and reference-free SV algorithms, splice detection tools, SNV/Indel callers and RNA abundance analysis, can capture virtually all somatic genomic alterations, providing both orthogonal validation and functional assessment of DNA mutations for precision medicine. The collation of large RNA-seq data sets has highlighted that paediatric cancer subtypes have unique gene-expression signatures^34^ and points towards the utility of RNA-seq as a valuable diagnostic tool in its own right. Together, RNA-seq provides an expanded capacity to identify alterations previously thought detectable only by WGS, raising the possibility that RNA-seq could be a stand-alone platform in precision medicine. However, the greatest capacity for precision comes from the integration of WGS and RNA-seq, since this currently provides the most precise and functional analysis, and as a consequence the greatest clinical impact.

## Methods

### Patient samples accrual

All patient samples were acquired from the Australian ZERO Childhood Cancer Precision Medicine Program as previously described in Wong *et al.*^11^. The data utilised in this study were obtained from ZERO through the pilot program TARGET which recruited patients between June 2015 to October 2017 and the PRISM clinical trial (NCT03336931) on patients collected between September 2017 and May 2021. Informed consent was collected for each patient enrolled. The TARGET trial was approved by the Sydney Children’s Hospitals Network Human Research Ethics Committee (LNR/14/SCH/497) and the PRISM trial was approved by the Hunter New England Human Research Ethics Committee of the Hunter New England Local

Health District (reference no. 17/02/015/4.06) and the New South Wales Human Research Ethics Committee (reference no. HREC/17/HNE/29). RNA-sequencing analysis and whole genome sequencing analysis presented in this paper consisted of all samples presented in Wong *et al.*^11^ and Lau *et al.*^13^ plus an additional 158 consecutive samples with matched WGS and RNA-seq data, resulting in a total cohort size of 477.

### Whole genome and RNA-sequencing analysis

Whole genome and whole transcriptome sample processing, library preparation, sequencing, alignment of raw fastq files, somatic variant analysis, tumour purity estimates, and structural variant calling was performed as previously described in Wong *et al.*^11^. In brief, RNA-seq reads were aligned using STAR (v2.5)^35^ two-pass method to the human genome assembly build hg19. Structural variants were identified using three methods: Arriba (v1.1.0)^7^, STAR-Fusion (v1.3.1)^8^ and JAFFA (v1.09)^9^.

### Reference-free structural variant and alternate splicing analysis

MINTIE is a reference-free SV detection method, previously demonstrated to excel at identifying intragenic SVs and aberrant splice variants^23^. We used MINTIE (v3.5)^23^ on 50 samples where all three rgRNA-seq methods missed the reportable SV that WGS had previously identified. We ran MINTIE without any controls, by setting parameter –p run_de_step=false. Results from MINTIE were filtered against the SVs identified from the matched WGS sample, all other results from MINTIE output were not explored further. Computation of MINTIE was performed on the National Computing Infrastructure and Pawsey Supercomputing Research Centre.

### RNA-seq Quantification and Expression Analysis

Aligned RNA-sequencing reads were quantified using RSEM (v1.2.31)^36^ to generate raw gene counts and transcripts per kilobase million (TPM). RNA abundance analysis was used to support the pathogenicity assessment of a SNV/Indel or SV. For each gene of interest, we identified a set of wildtype controls from the cohort, that had no detectable SNV/Indel, CNV or SV. From these wildtype controls, z-scores and fold changes can be calculated using standard methods.

### RNA-sequencing-based SNV/Indel variant calling

SNVs were called on RNA-seq data using GATK HaplotypeCaller (v3.6) and Indels were called using RNAIndel^37^, both using default settings. Variants were filtered to only the somatic variants identified from the WGS data, and then to those occurring within coding regions or splice-altering intronic variants. The variant allele frequency (VAF) was then calculated on the RNA-seq mutations and compared to the WGS VAF to determine the zygosity state of each mutation relative to the DNA.

### Carbonite a next-flow pipeline for RNA-seq analysis

From our findings in this paper we developed *Carbonite*, an open-source scalable RNA-seq analysis pipeline for cancer precision medicine. This pipeline uses STAR^35^ for alignment, and RSEM^36^ for quantification. It incorporates two of our rgRNA-seq fusion detection algorithms (STAR-Fusion^8^ and Arriba^7^). Due to long run-times and compute usage JAFFA^9^ needs to be run independently. It also incorporates rfRNA-seq SV and splice detection tool MINTIE^23^. After duplicates marked it runs SNV/Indel detection using GATK Haplotype caller and RNAIndel^37^. There are additional tools included in this workflow which are outside the scope of this paper. This pipeline can be installed using the command `git clone` git@github.com::<u>CCICB/nf-carbonite.git</u>.

### Statistics

All statistical analysis and visualisations were performed in R (v3.4.0).

## Data availability

The data generated in this study are from the ZERO Childhood Cancer Program. All patients have provided consent for secondary research use. Summary molecular and basic clinical data (e.g. diagnosis, stage, gender, age) are publicly accessible at https://beacon.zerochildhoodcancer.cloud/ which uses the GA4GH Beacon v2 protocol. Privacy-sensitive, patient-level whole-genome and whole-transcriptome data (including raw fastq and BAM files and processed variant calls from WGS and RNA-seq) are available through ZERO’s Data Access Committee. Full details are provided at https://www.zerochildhoodcancer.org.au/clinicians-researchers/for-researchers/data-and-sample-resources.

## Code availability

The pipeline called *Carbonite* and its associated scripts developed from this paper has been turned into a Nextflow pipeline and is available at https://github.com/CCICB/nf-carbonite for use with either the hg19 or hg38 human genome reference.

## Acknowledgments

We sincerely thank patients and parents for participating in this study. We thank the many clinicians, tumour banks and health professionals for their time acquiring consent for patients and for the collection and coordination of samples and associated clinical data at Sydney Children’s Hospital, Randwick; the Children’s Hospital at Westmead; the John Hunter Children’s Hospital; the Queensland Children’s Hospital; the Royal Children’s Hospital Melbourne; the Monash Children’s Hospital; the Adelaide Women & Children’s Hospital; and the Perth Children’s Hospital. Tumour samples and coded data were supplied by the Children’s Cancer Centre Biobank at the Murdoch Children’s Research Institute and The Royal Children’s Hospital (https://mcri.edu.au/research/projects/childrens-cancer-centre-biobank). The establishment and running of the Children’s Cancer Centre were made possible through generous support by Cancer In Kids @ RCH (www.cika.org.au), the Royal Children’s Hospital Foundation and the Murdoch Children’s Research Institute. We thank ANZCHOG as the trial sponsor, and we thank the staff of the Clinical Translation Theme of the Children’s Cancer Institute for their dedicated work on the ZERO Childhood Cancer Program. We thank the Australian BioCommons, Velsera/SevenBridges and Children’s Hospital of Philadelphia Center for Data-Driven Discovery in Biomedicine for supporting the development of the CAVATICA genome analysis platform. The authors acknowledge the provision of computing and data resources provided by the Australian BioCommons Leadership Share (ABLeS) program. This program is co-funded by Bioplatforms Australia (enabled by NCRIS), the National Computational Infrastructure and Pawsey Supercomputing Research Centre. ZERO Childhood Cancer is a joint initiative led by Children’s Cancer Institute and the Kids Cancer Centre, Sydney Children’s Hospital, Randwick. The authors would like to acknowledge Luminesce Alliance—Innovation for Children’s Health for its contribution and support. Luminesce Alliance is a not-for-profit cooperative joint venture between the Sydney Children’s Hospitals Network, the Children’s Medical Research Institute and the Children’s Cancer Institute. It has been established with the support of the NSW Government to coordinate and integrate paediatric research. Luminesce Alliance is also affiliated with the University of Sydney and the University of New South Wales Sydney. This work has been funded by the Australian Federal Government Department of Health, the New South Wales State Government and the Australian Cancer Research Foundation for funding to establish infrastructure to support the ZERO Childhood Cancer personalised medicine programme. Funding from the Kids Cancer Alliance, Cancer Therapeutics Cooperative Research Centre supported the development of a personalised medicine programme; Tour de Cure supported tumour biobank personnel; the Steven Walter Children’s Cancer Foundation and the Hyundai Help 4 Kids Foundation supported G.M.M. and P.G.E.; Samuel Nissen Charitable Foundation supported P.G.E.; the Lions Kids Cancer Genome Project is a joint initiative of Lions International Foundation, the Australian Lions Children’s Cancer Research Foundation (ALCCRF), the Garvan Institute of Medical Research, the Children’s Cancer Institute and the Kids Cancer Centre, Sydney Children’s Hospital. Lions International and ALCCRF provided funding to perform WGS and for key personnel, with thanks to J. Collins for project governance and advocacy. The Cure Brain Cancer Foundation supported the RNA sequencing of patients with brain tumours; the Kids Cancer Project supported molecular profiling and molecular and clinical trial personnel; and the University of New South Wales, W. Peters and the Australian Genomics Health Alliance provided personnel funding support. The Medical Research Future Fund, Australian Brain Cancer Mission/National Health & Medical Research Council/Lifting Clinical Trials and Registry Capacity (NHMRC MRF9500002), the Minderoo Foundation’s Collaborate Against Cancer Initiative and funds raised through the ZERO Childhood Cancer Capacity Campaign, a joint initiative of Children’s Cancer Institute and the Sydney Children’s Hospital Foundation, supported the national clinical trial and associated clinical and research personnel. Cancer Institute of New South Wales and New South Wales Health (fellowship funding for M.J.C.; CINSW Program Grant 2019/TPG2037 for M.J.C and D.S.Z and 2021/TPG2112 for G.M.M., M.H., and L.M.S.L). National Health and Medical Research Council (Synergy Grants APP2018642 for G.M.M, M.H., and L.M.S.L., and APP2019056 for D.S.Z; Leadership Grant APP2017898 for D.S.Z.). This research was supported by an Australian Government Research Training Program (RTP) Scholarship (for C.M.). The 2018 Priority-Driven Collaborative Cancer Research Scheme, co-funded by Cancer Australia and My Room, for personnel and computational support (APP1165556 awarded to M.J.C.). J.R.H is supported by the Hospital Research Foundation and McClurg Foundation. R.E. has support from a Cancer Council of Western Australia Research Fellowship and a Brainchild Fellowship from the Pirate Ship Foundation. N.G.G is funded by the Perth Children’s Hospital Foundation Stan Perron Chair in Paediatric Oncology and Haematology.

## Extended Data Figures

**Extended Data Figure 1.**
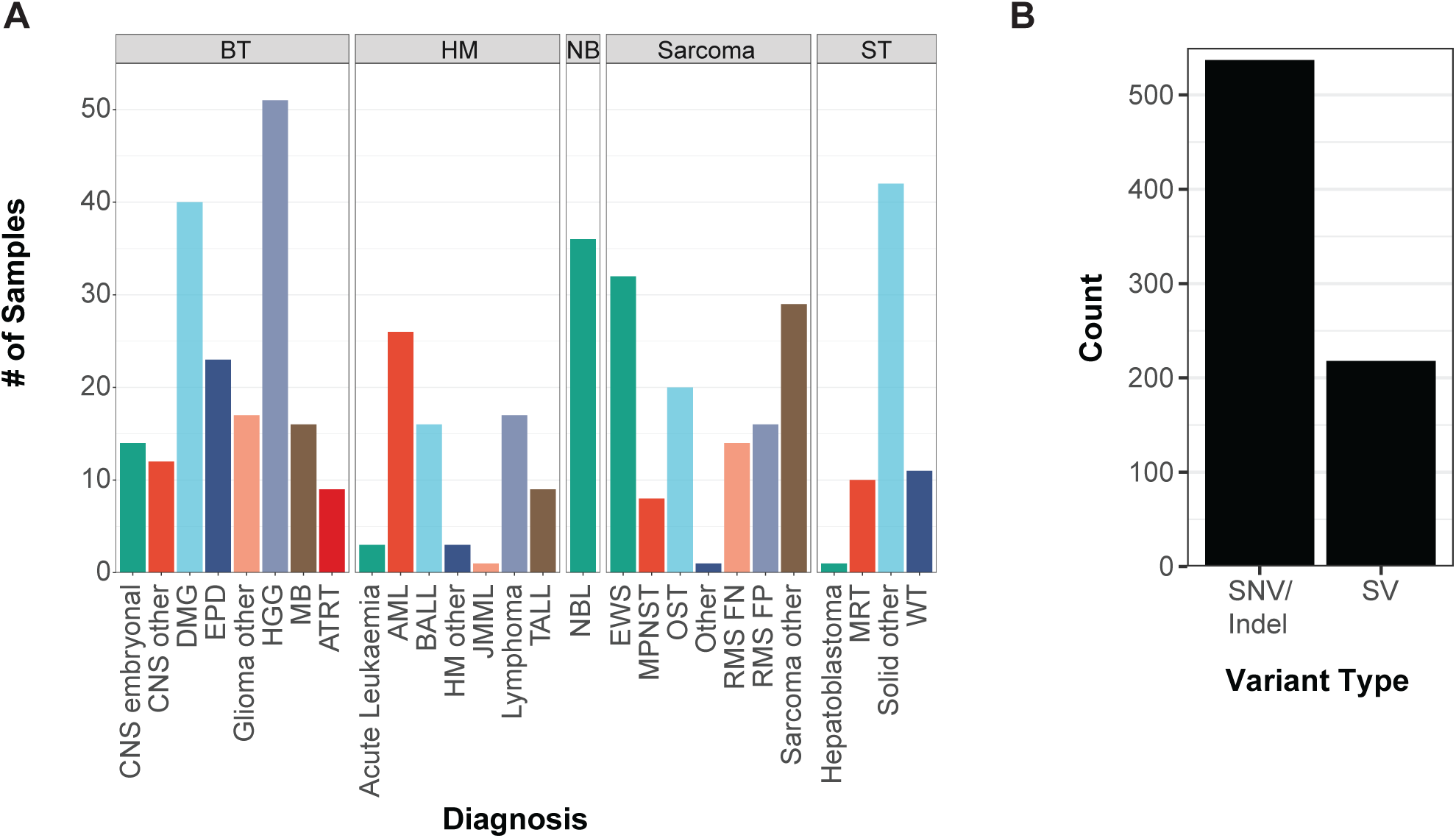
Overview of the paediatric cancer cohort. (A) Composition of the cohort comprising 477 high-risk paediatric cancer samples profiled by both WGS and RNA-seq. Samples are grouped into 5 major diagnostic categories: brain tumours (BT), haematological malignancies (HM), neuroblastomas (NB), sarcomas, and other solid tumours (ST). **(B)** The number of reportable variants, defined as likely pathogenic or pathogenic, identified across the cohort, stratified by variant type: single nucleotide variants and small insertions/deletions (SNV/Indels), and structural variants (SV).

**Extended Data Figure 2.**
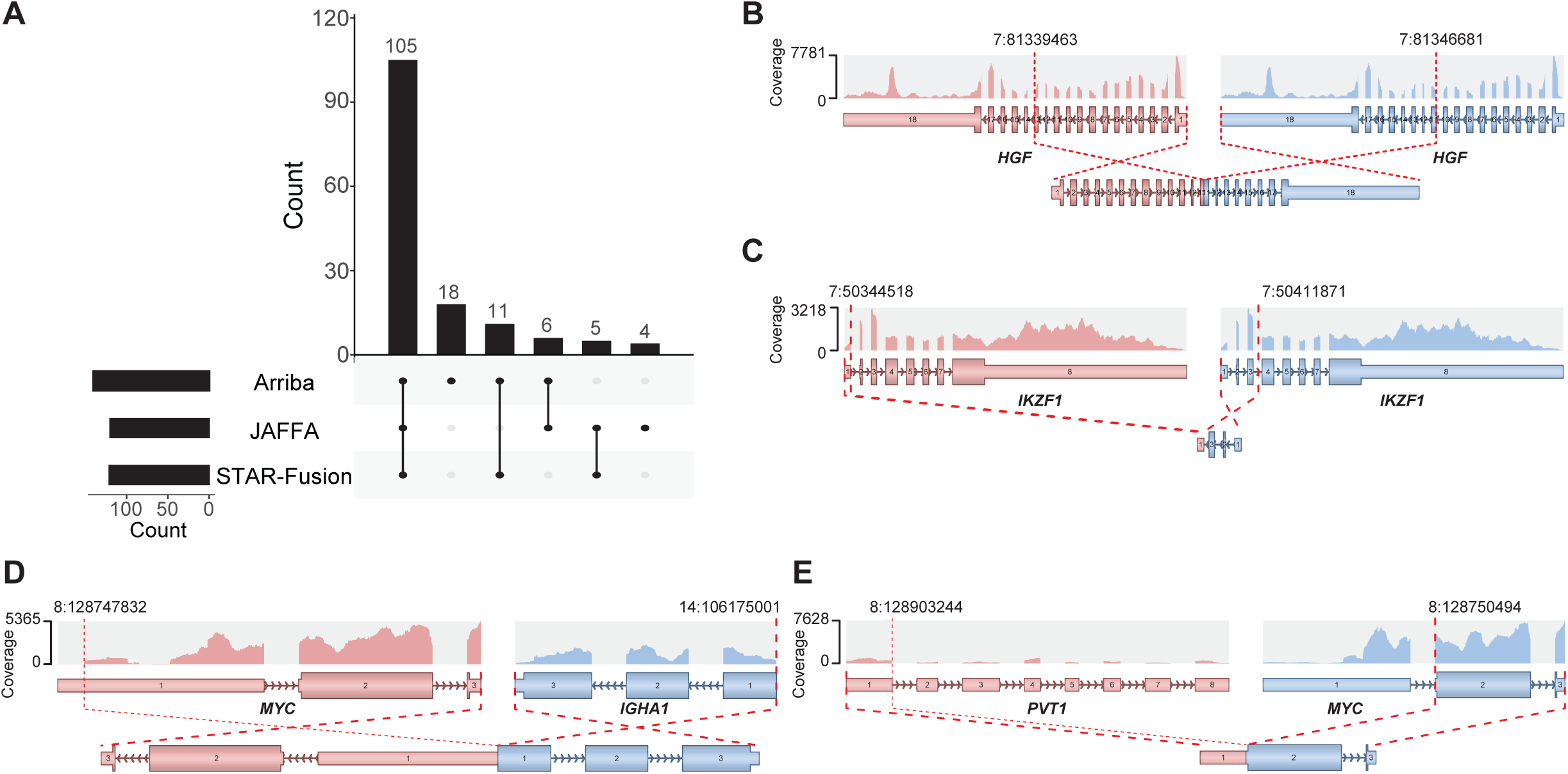
SVs identified by reference-guided RNA-seq algorithms. (A) UpSet plot showing the intersection of gene fusions detected by three reference-guided algorithms: Arriba, JAFFA and STAR-Fusion used in our initial RNA-seq pipeline. Arriba plots showing the transcribed exons, expression coverage, and breakpoint locations for selected SVs, including an **(B)** *HGF* internal tandem duplication of exons 11-13, **(C)** *IKZF1* inversion, **(D)** *MYC::IGHA1* fusion and **(E)** *PVT1::MYC* fusion. Panels B and C are examples of SVs uniquely identified using Arriba, undetected by JAFFA and STAR-Fusion. Panels D and E illustrate enhancer hijacking events involving *MYC*, where breakpoints occurred within the transcribed region of *MYC*, and thus captured by rgRNA-seq.

**Extended Data Figure 3.**
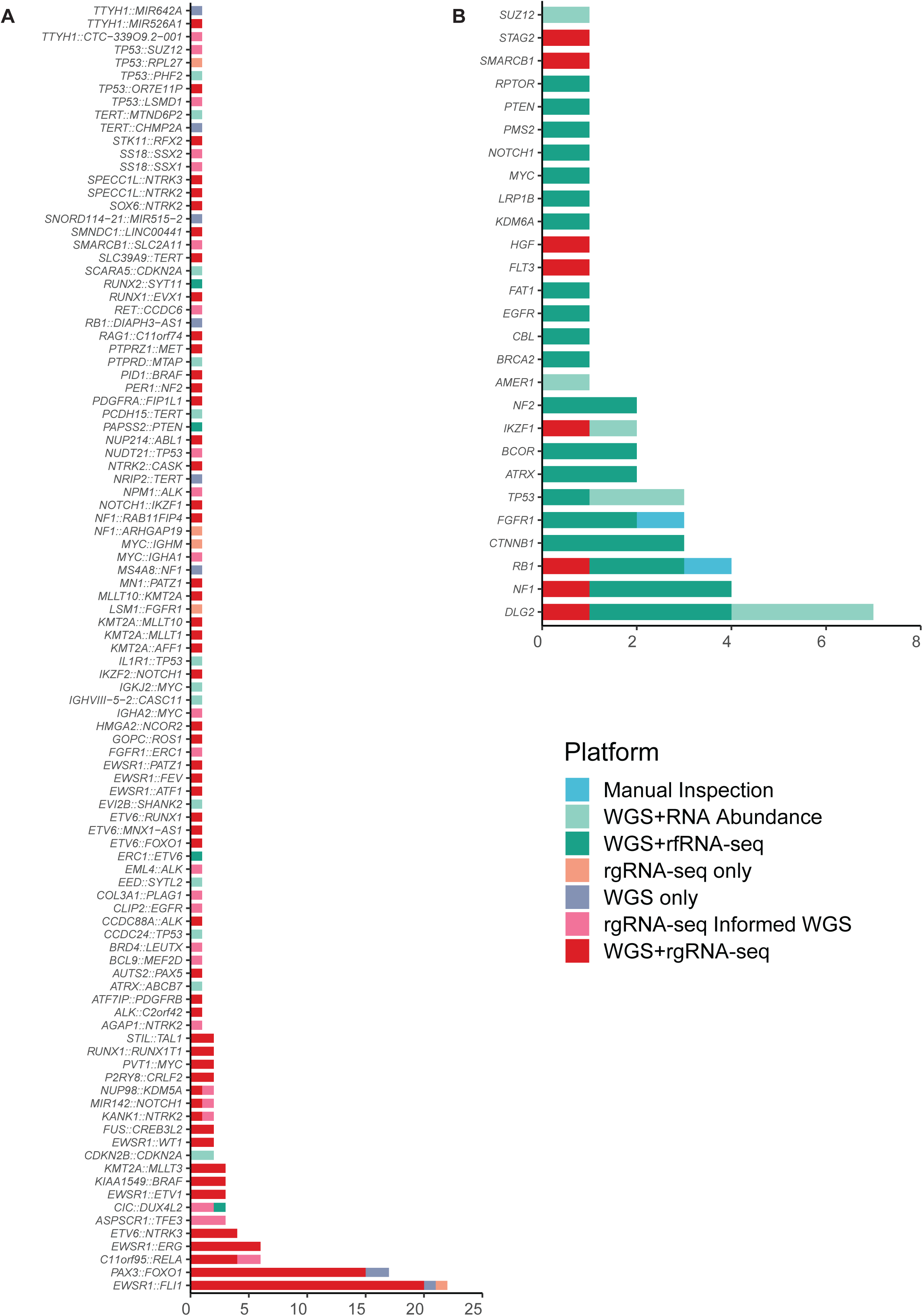
SV detection by platform and methodology The frequency of. (A) gene fusions and **(B)** intragenic SVs identified by different platforms or approaches: WGS only (purple), rgRNA-seq only (peach), both (red), initially detected by rgRNA-seq and later identified in WGS via LINX (pink), WGS and rfRNA-seq (dark green), WGS with RNA abundance support (teal), or through manual inspection using Integrative Genome Viewer (IGV; blue).

**Extended Data Figure 4.**
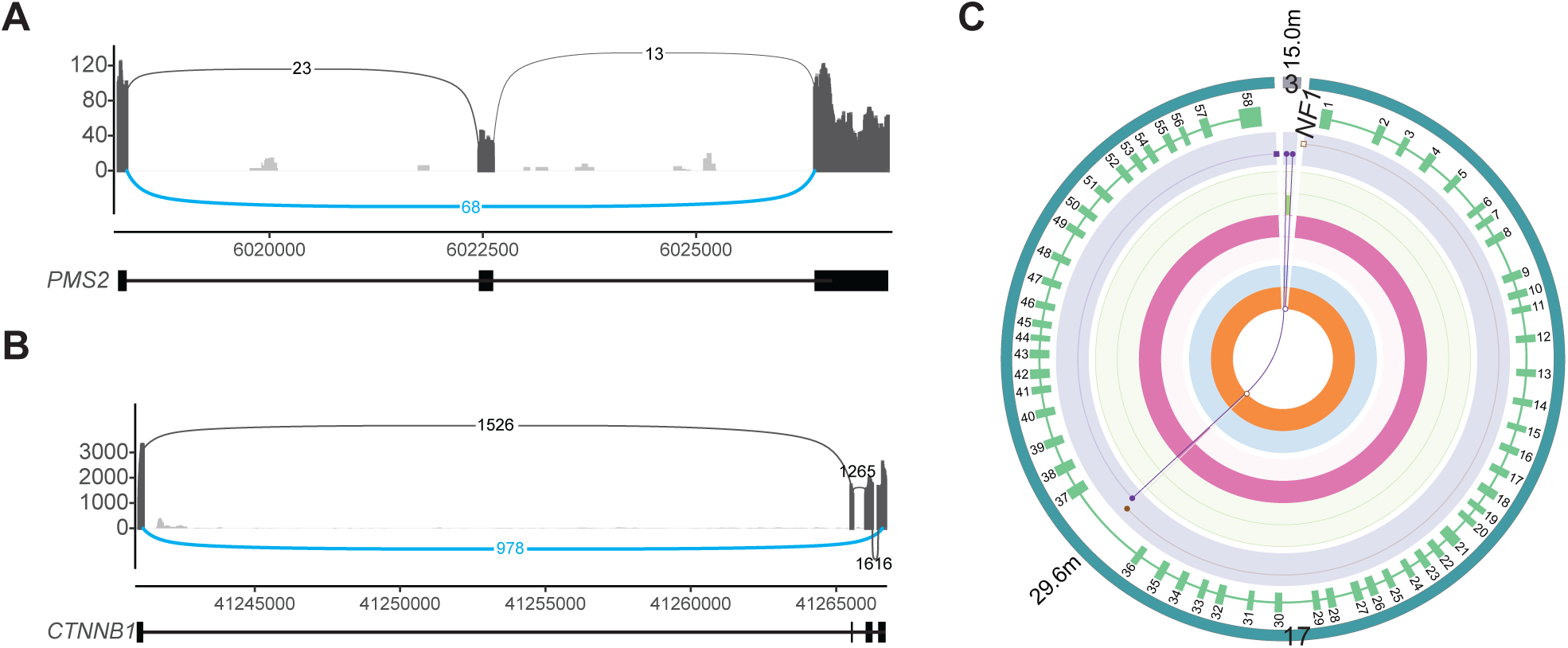
Intragenic SVs identified using reference-free RNA-seq. Sashimi plots showing aberrant splicing due to intragenic deletions identified by reference-free RNA-seq in **(A)** *PMS2* and **(B)** *CTNNB1*. **(C)** LINX plot showing the intragenic insertion of 17 nucleotides from chromosome 3 into intron 36 of *NF1*, an event not detected by rgRNA-seq but identified using the rfRNA-seq approach.

## Supplementary Information

Supplementary Table 1. Variant Allele Frequencies of reportable SNVs and Indels in WGS and RNA-seq.

